# Rapid environmental change favours the evolution of shorter lifespan

**DOI:** 10.1101/2020.01.29.924373

**Authors:** Martin I. Lind, Irja I. Ratikainen, Johan Andersson, Hanne Carlsson, Therese Krieg, Tuuli Larva, Alexei A. Maklakov

## Abstract

The role of environmental variation in the evolution of lifespan is contested. Classic theory predicts that variable environments favour the evolution of long life, but recent theoretical work suggests that environmental variation can instead select for shorter lifespan when the changes are rapid relative to generation time. Here, we used experimental evolution to study how lifespan evolves in response to temperature variation. Genetically diverse populations of the outcrossing nematode *Caenorhabditis remanei*, were exposed to a novel, stressful temperature for 30 generations, under stable, gradually warming or fluctuating thermal regimes. We found the evolution of shorter lifespan in populations exposed to environments that fluctuated within the individual’s lifetime, compared to populations evolving in stable warm temperature. While climate warming is predicted to increase environmental stochasticity, our results show that fast temperature cycles can rapidly select for short lifespan.

## Introduction

Natural environments are rarely constant. Instead, they change directionally or fluctuate across timescales, and such variation can strongly influence evolution of life-histories. Accordingly, understanding how the organisms adapt to environmental heterogeneity has long been a central question in evolutionary biology (Murphy, 1968; Tuljapurkar & Orzack, 1980; Charlesworth, 1994; Roff, 2002; Wilbur & Rudolf, 2006; Morris *et al*., 2008; Cotto & Ronce, 2014; Botero *et al*., 2015; Tufto, 2015; Ratikainen & Kokko, 2019).

Evolution in novel and variable environments can have far-reaching consequences for the evolution of life-history strategies, especially for the trade-off between growth, reproduction and somatic maintenance, which directly affects the evolution of lifespan. The theoretical models often make opposing predictions, from the evolution of robust and long-lived individuals, to the evolution of short lifespan in heterogeneous environments (Murphy, 1968; Tuljapurkar & Orzack, 1980; Charlesworth, 1994; Wilbur & Rudolf, 2006; Morris *et al*., 2008; Cotto & Ronce, 2014; Ratikainen & Kokko, 2019).

Evolution of lifespan in novel but stable environments is generally considered as changes in age-specific mortality (Williams *et al*., 2006), where increased random adult mortality in the new environment is predicted to result in short lifespan, which has also been demonstrated empirically by experimentally elevating mortality rates but keeping the environment constant (Stearns *et al*., 2000). However, novel environments may also supress reproduction, which can strengthen selection for somatic maintenance and result in the evolution of longer lifespan by increasing the value of survival (Irish *et al*., 2025). In addition, mortality in the novel environment is seldom random but can be condition dependent and select upon specific phenotypes. Condition dependent mortality have been empirically shown to result in the evolution of stress resistance and long lifespan (Reznick *et al*., 2004; Chen & Maklakov, 2012). Finally, empirical studies show that selection in stressful but stable environments often results in the evolution of stress resistance and longer lifespan but trade-offs with growth or reproduction (Sørensen *et al*., 2003; Bubliy & Loeschcke, 2005; Chen *et al*., 2016; Lind *et al*., 2017).

Traditionally, variable environments were also expected to favour long lifespan, because individuals that survive unfavourable periods may reproduce later when conditions improve (Murphy, 1968; Tuljapurkar & Orzack, 1980; Charlesworth, 1994; Wilbur & Rudolf, 2006; Morris *et al*., 2008). More recent theory, however, shows that this prediction depends critically on the tempo of environmental change relative to generation time and on whether organisms can adjust their phenotype after development. (Cotto & Ronce, 2014; Ratikainen & Kokko, 2019). Central to the new theories is the concept of phenotypic mismatch. If phenotypes are developmentally stable or plasticity is irreversible (developmental plasticity), phenotypic mismatch can select for the evolution of short lifespan in order to complete most of reproduction in the environment of development (Cotto & Ronce, 2014; Ratikainen & Kokko, 2019). Alternatively, phenotypic mismatch can select for the evolution of reversible plasticity to track environmental changes, accompanied by the evolution of long lifespan (Ratikainen & Kokko, 2019). Without reversible plasticity, stable but novel as well as heterogeneous environments should select for short lifespan (Cotto & Ronce, 2014).

Since adaptation typically proceeds with a lag, an adapting population is not instantly at the new fitness peak (Lande & Shannon, 1996; Kopp & Matuszewski, 2014). Because selection gradients on traits decline with age (Medawar, 1952; Hamilton, 1966; Caswell & Shyu, 2017), older individuals have a weaker response to selection and are thus further from the new fitness peak. Therefore, in a population with age structure, adaptation to new environments is associated with reduced fitness, which is most severe for late age classes, resulting in weaker selection on alleles with late-acting deleterious effects. While a rapid shift to a new stable environment only results in a transient effect, a population in a continuously moving or fluctuating environment will never be fully adapted, and selection against somatic maintenance strongest (Cotto & Ronce, 2014). Taken together, these ideas generate a clear prediction, namely, when environmental change is rapid relative to generation time, short lifespan should be favoured unless reversible plasticity can effectively reduce mismatch. These predictions are especially relevant in the context of climate change. Ongoing warming is expected not only to increase mean temperatures, but also to increase environmental variability and stochasticity. Despite this, empirical tests of how the timescale of environmental change shapes lifespan evolution remain scarce.

We set out to test these predictions using an experimental evolution approach in the dioecious nematode *Caenorhabditis remanei*, adapting to five different temperature regimes (figure 1). Genetically heterogeneous populations, originally adapted to 20°C, evolved for approximately 30 generations under five thermal regimes: constant 20°C, constant 25°C, gradual warming from 20°C to 25°C, and fluctuations between 20°C and 25°C occurring either every second or every tenth generation. Because the experiment used overlapping generations, it directly captures the temporal structure central to recent theory. We then asked how these thermal regimes affected the evolution of lifespan and age-specific mortality. Based on theory, we predicted that rapid temperature cycles would favour shorter lifespan, whereas slower or more stable regimes would be less likely to do so.

**Figure 1.**
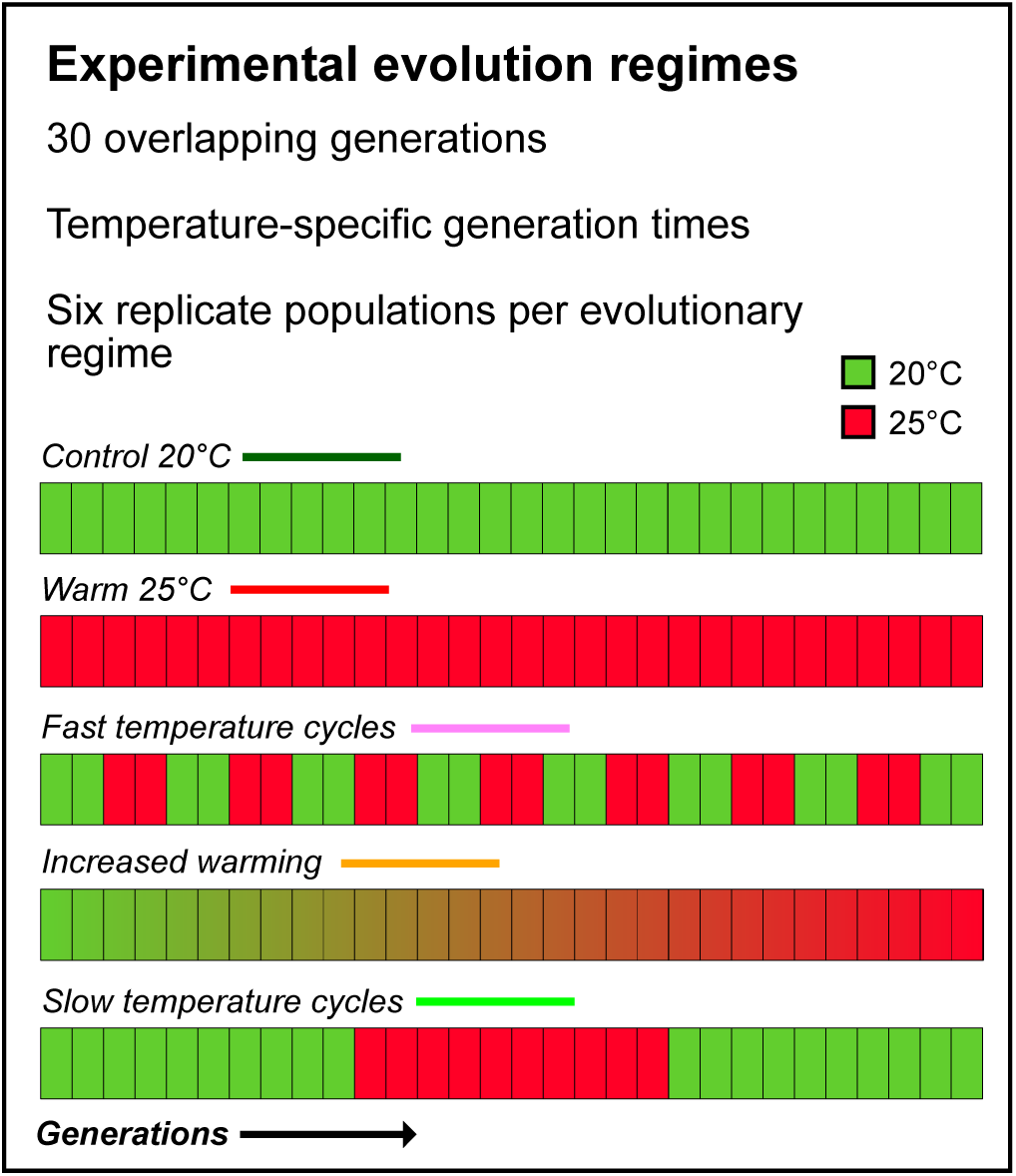
Overview of the experimental evolution regimes. Each of the five regimes consisted of six replicate populations. Generation time was 4 days in 20°C and 3.4 days in 25°C, and we used an overlapping generations setting for approximately 30 generations.

## Methods

### Experimental evolution

Experimental evolution was conducted with the SP8 strain of *C. remanei* as founder population (obtained from N. Timmermeyer, then at the Department of Biology, University of Tübingen, Germany). This strain harbours substantial standing genetic variation for life-history traits and responds readily to selection (Chen and Maklakov 2012, 2014; Lind et al. 2017). The strain has been lab-adapted for 15 generations at 20°C prior to the setup of the experimental evolution.

The experimental evolution procedure is described in detail in Lind *et al*., (2020) and outlined in figure 1. We conducted experimental evolution for approximately 30 generations using three different temperature settings: 20°C, 25°C and slow increase from 20°C to 25°C by 0.1°C every 2 days. Importantly, by generation we refer to generation time, as we used an overlapping generations setting. Thus, the actual number of generations in each regime may deviate from 30 generations, if generation time itself evolves. We set up five selection regimes. *Control 20°C* populations were experiencing 20°C for 30 generations, and *Warm 25°C* populations were experiencing 25°C for 30 generations. *Increased warming* populations experienced slow increase from 20°C at the start of the experiment to 25°C on the last day of selection in slow incremental steps of 0.1°C over the period of 30 generations. In *Slow temperature cycles,* populations were experiencing the first ten generations from the start of the experiment in 20°C, then were switched for the next ten generations to 25°C, and then switched back again for the remaining ten generations to 20°C. Finally, the *Fast temperature cycles* regime spent the first two generations at the start of the experiment in 20°C, then were switched for two generations to 25°C and this cycle was repeated 7.5 times (the last two generations were in 20°C).

We defined generation time as the average difference in age between parents and offspring (Charlesworth, 1994) in 20°C and 25°C, and it was calculated by solving the Euler-Lotka equation from the life-table of age-specific survival and reproduction (Stearns, 1992; Charlesworth, 1994) using trial data from the original SP8 line.

Generation time was 4.0 days in 20°C and 3.4 days in 25°C, which resulted in 120 days of selection for *Control 20°C*, 114 days for *Slow temperature cycles*, 110 days for *Increased warming* and *Fast temperature cycles* and finally 101 days for *Warm 25°C*. To ensure equal exposure to the two temperature treatments during biological time, the worms spent shorter chronological time in 25°C than in 20°C for the two temperature cycle treatments, because generation time is faster in 25°C.

During experimental evolution, the populations were kept on 92 mm NGM-plates (Stiernagle, 2006). The antibiotics kanamycin and streptomycin, and the fungicide nystatin was added to agar and LB to combat bacterial and fungal infections (Lionaki & Tavernarakis, 2013; Lind *et al*., 2016). We seeded the plates with the antibiotic-resistant *E. coli* strain OP50-1 (pUC4K) from J. Ewbank at Centre d’Immunologie de Marseille-Luminy, France. The lines were kept in exponential growth phase by cutting out a bit of agar containing 150 individuals of mixed ages and transferring this to freshly seeded plates, in order to keep the populations age-structured in overlapping generations. Transfer was conducted when needed (every one to two days) and plates were never allowed to run out of food. We set up six independent replicate populations of each selection treatment, resulting in 30 experimental populations in total. All populations were expanded in size for two generations and frozen after 10, 20 and 30 generations. For generations 10 and 20, a random subset of each population was used for freezing, to not interfere with the ongoing experiment.

### Lifespan assays

Before assays, worms were recovered from freezing and grown for two generations in common garden at 20°C, each generation synchronized by bleaching, a standard procedure that kills all life-stages but the eggs (Stiernagle, 2006). Lifespan assays were established using ten age-synchronised females in the L4 stage as target worms, and to keep them mated they were kept together with ten background males from the original SP8 line. Worms were developing in the testing temperature since eggs.

Worms were transferred daily, and sex ratio was kept at 1:1 by matching the number of background males to the number of testing females alive. The assay was run using four replicate plates of each line and temperature (20°C and 25°C) combination, resulting in 240 plates (five selection regimes × six replicate lines × two temperatures × four replicate plates) and 2400 target worms. To separate the effect of temperature and cabinet, two cabinets were used for each temperature, and 2 replicate plates of each line was run in each cabinet. Worms were scored as dead if they did not respond to gentle prodding by the picker, and dead by matricide if death was likely caused by bagging. Missing worms were censored.

### Statistical analyses

Survival was analysed using Cox proportional hazard models with Gaussian random effects implemented in the *coxme* package for *R 4.32*. Selection treatment and testing temperature were fitted as fixed factors, and population, plate and testing cabinet (climate chamber) as random effects. Significance of main effects were evaluated using the *car* package and planned contrast were performed using the *emmeans* package. Of the 2400 worms, 18 were removed because of developmental abnormalities or loss of the whole plate during handling, and therefore not included in analyses. The number of worms in the models were as follows*. Control 20°C*: n = 233 in 20°C, n = 240 in 25°C. *Fast temp. cycles*: n = 239 in 20°C, n = 240 in 25°C. *Increased warming*: n = 234 in 20°C, n = 240 in 25°C. *Slow temp. cycles*: n = 239 in 20°C, n = 240 in 25°C. *Warm 25°C*: n = 237 in 20°C, n = 240 in 25°C.

Mortality rate was analysed in a Bayesian framework using the *BaSTA 2.0.1* package (Colchero *et al*., 2012), which performs a capture-mark-recapture (CMR) analysis and is therefore well suited to deal with censored data. To find the best model, we fitted mortality rates using Gompertz, Weibull and logistic functions, with simple, Makeham or bathtub shapes, and these models were compared using DIC (the Bayesian equivalent to AIC). Since models with estimated intercept did not converge, we fitted the *Control* treatment as the baseline. That enabled all other experimental evolution treatments to be compared against each other. All models were run as four independent simulations; each simulation was run for 600.000 iterations, with the first 6.000 iterations discarded as burn in, and a model sample was taken every 600 iteration. We found that the most appropriate mortality function (lowest DIC) was the Weibull model with a Makeham shape. The three parameters of the mortality function describe how the mortality rate changes with age. Specifically, c represent the baseline (extrinsic) mortality, b_0_ the overall level of age-related mortality, and b_1_ the increase age-dependent mortality with increased age.

The difference between the posterior distributions of the selection regime and temperature parameters was assessed using the Kullback-Lieber divergence calibration (KLDC) (Kullback & Leibler, 1951; Karabatsos, 2006), where a value of 0.5 signifies identical distributions, while a value of 1 signifies completely non-overlapping distributions. KLDC values above 0.8 are generally considered indicative of differences between compared distributions (Rodríguez-Muñoz *et al*., 2019; Spagopoulou *et al*., 2020).

## Results

### Lifespan

No experimental evolution regime × temperature interaction was detected (χ^2^ = 1.92, df = 4, p = 0.751), implying that the ordering of the evolution regimes was not different in the two temperatures. This interaction was therefore removed from the final model. We found strong evidence that female lifespan was shorter in 25°C than in 20°C (χ^2^ = 496.12, df = 1, p < 0.001), and evidence for lifespan differences between the experimental evolution regimes (χ^2^ = 9.97, df = 4, p = 0.041, Figure 2, Table 1, supplementary figure 1). Planned post-hoc contrasts showed that the *Fast temperature cycle* regime had evolved a shorter lifespan than the long-lived *Warm 25°C regime* (z = -3.043, p = 0.009) but did not significantly differ in lifespan from the other regimes (*Increased warming* regime: z = -2.219, p = 0.090; *Slow temperature cycle:* z = -1.565, p = 0.334; *Control 20°C:* z = -1.752, p = 0.241, supplementary table 1). This contrast was also significant when performing all possible post-hoc comparisons, even if they were not planned a priori (supplementary table 2).

**Figure 2.**
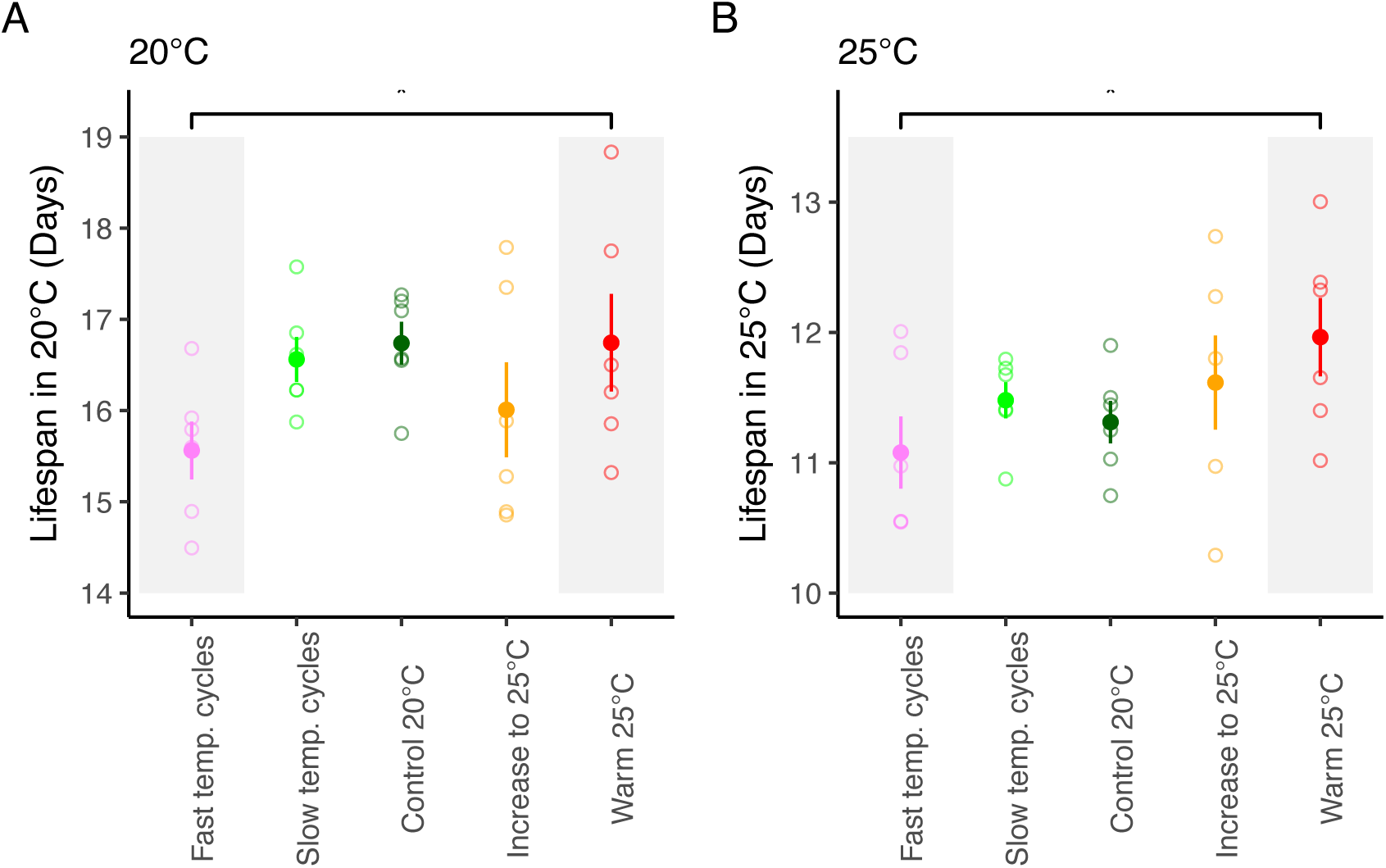
Mean lifespan in (A) 20°C and (B) 25°C. Symbols represent experimental evolution regime (mean ± SE based upon population means). Open symbols represent the mean of each replicate population. The grey boxes represent the groups where we found evidence for differences in lifespan across both temperatures, no temperature interaction was detected.

**Table 1.**
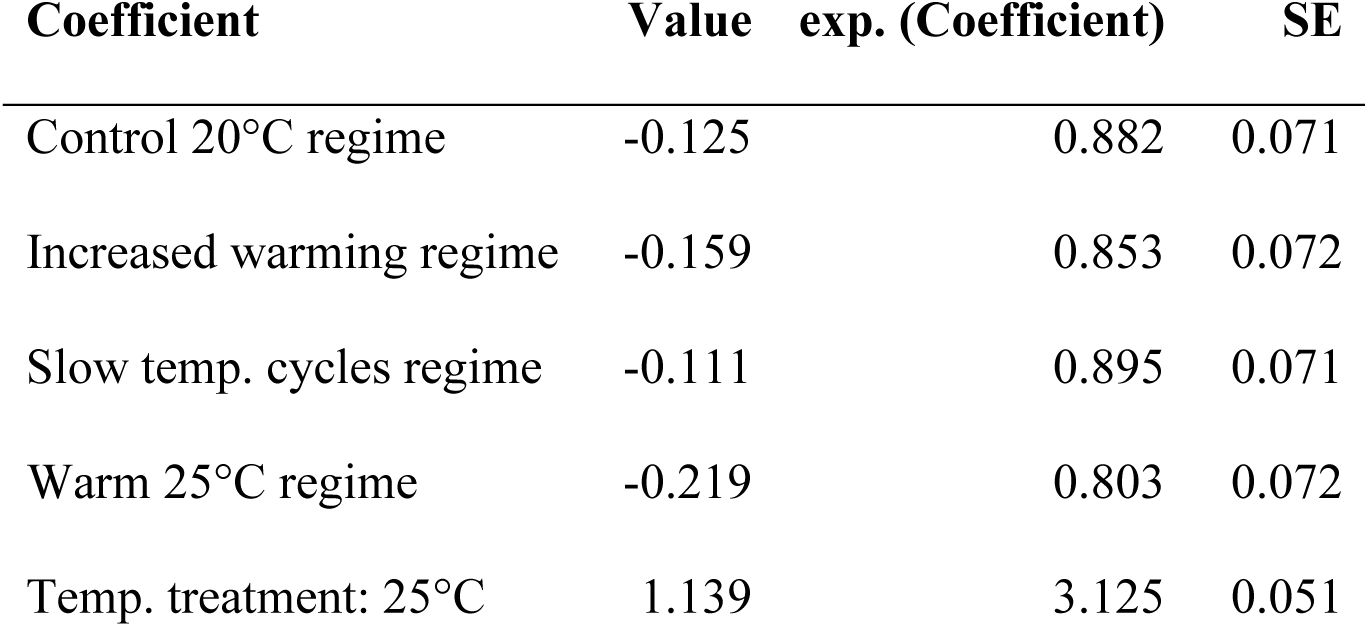
Coefficients from the best model of the survival analysis, which does not involve the non-significant interaction between Regime and Temperature. All comparisons in the table are made between the *Fast temperature cycle* regime and the temperature treatment 20°C.

| Coefficient | Value | exp. (Coefficient) | SE |
| --- | --- | --- | --- |
| Control 20°C regime | -0.125 | 0.882 | 0.071 |
| Increased warming regime | -0.159 | 0.853 | 0.072 |
| Slow temp. cycles regime | -0.111 | 0.895 | 0.071 |
| Warm 25°C regime | -0.219 | 0.803 | 0.072 |
| Temp. treatment: 25°C | 1.139 | 3.125 | 0.051 |

### Mortality rate

The most appropriate mortality function with lowest DIC was a Weibull model with Makeham structure.

Mortality patterns were strongly influenced by both temperature and selection regime (figure 3). KLDC comparisons between 25°C and 20°C revealed near-complete divergence across all mortality parameters (c, b_0_, b_1_; KLDC ≥ 0.95), indicating that higher temperature substantially increased both age-independent mortality and age-dependent mortality, affecting both background mortality and the rate of increase with age (figure 4).

**Figure 3.**
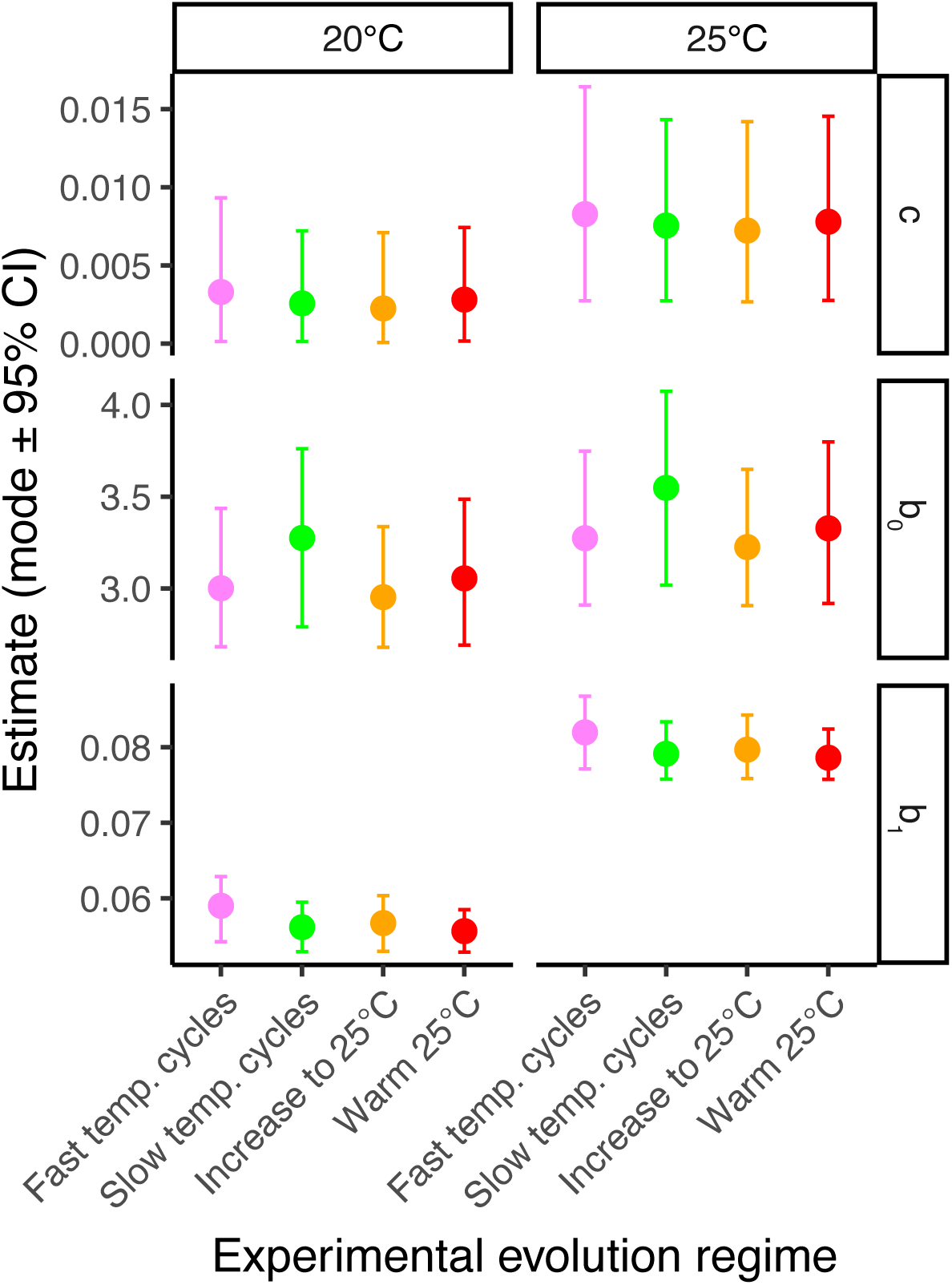
Mortality parameter estimates (posterior mode ± credible interval) based on a Weibull model with Makeham structure. c represents the baseline (extrinsic) mortality, b_0_ the overall level of age-related mortality, and b_1_ the increase age-dependent mortality with increased age.

**Figure 4.**
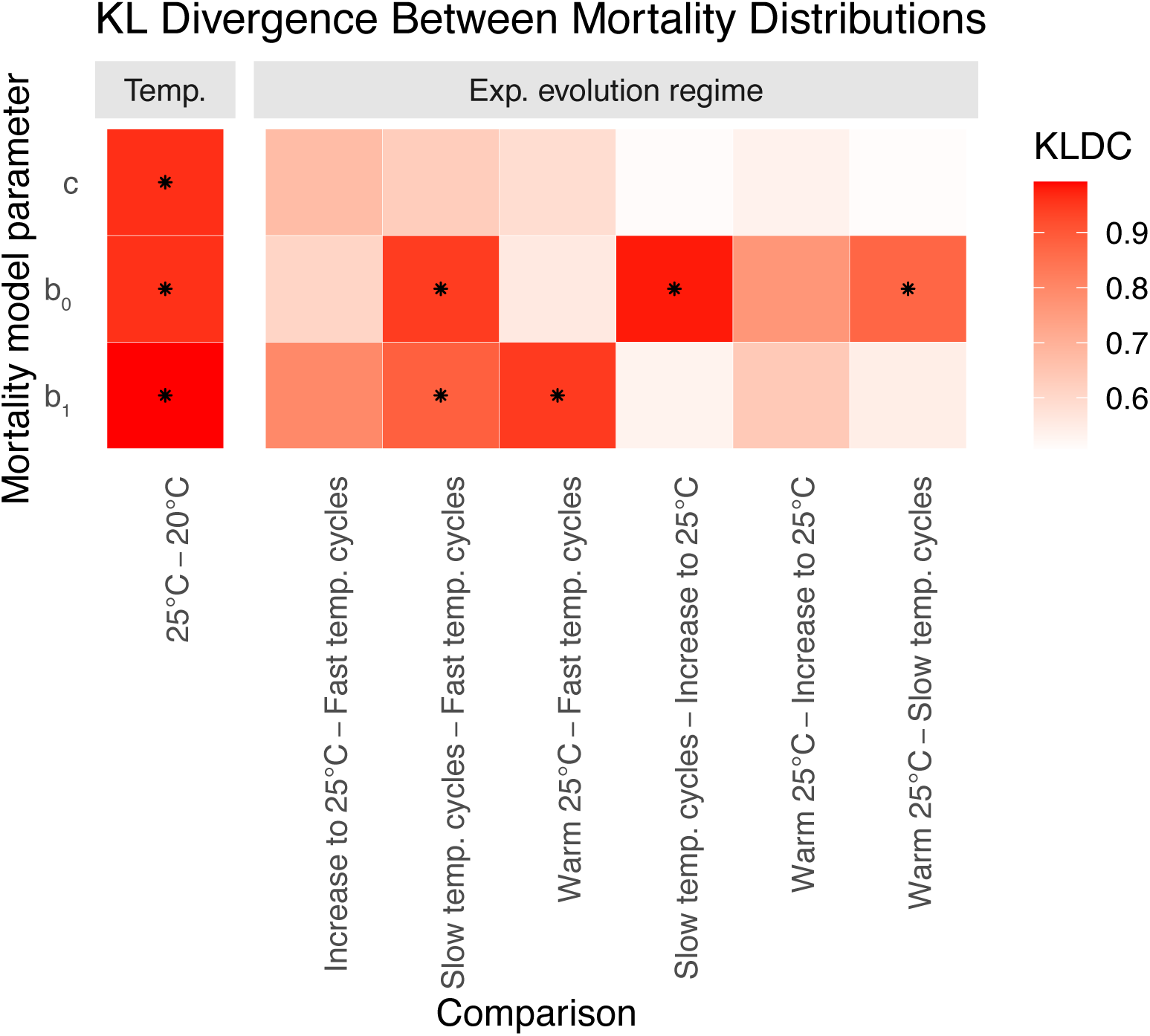
Heatmap of KLDC divergence between mortality parameters (c, b_0_ and b_1_) for temperature and for the comparisons of the experimental evolution regimes. KLDC > 0.8 indicate statistically different distributions and are visualised by asteriscs (*). Exact KLDC values are presented in supplementary table 3.

Using the *Fast temp. cycles* regime as the reference, the KLDC analysis also highlighted clear differences among selection lines (figure 3-4, supplementary table 3). The *Warm 25°C* regime differed predominantly in the increase of age-dependent mortality rate with age (b_1_ = 0.95), with moderate divergence in the overall mortality rate (b_0_ = 0.56) and background mortality (c = 0.59). Also the *Slow temperature cycle* regimes had strong differences in both baseline (b_0_ = 0.95) and age-dependent mortality rate (b_1_ = 0.97) relative to the *Fast temperature cycle* regime, while the divergence in age-independent mortality was moderate. In contrast, the *Increased warming regime* differed only moderately from the *Fast temperature cycle regime*, with all mortality parameters with KLDC < 0.8. Pairwise comparisons among the other regimes (excluding the comparisons with the baseline *Fast temperature cycles*) revealed moderate-to-strong divergence, particularly in age-dependent baseline mortality parameters (b_0_), indicating variation in mortality patterns across the evolutionary regimes (Figure 4).

## Discussion

Theory makes opposing predictions about how environmental heterogeneity shapes life histories, ranging from robust, slow-reproducing, long-lived phenotypes to fragile, fast-reproducing, short-lived ones (Murphy, 1968; Tuljapurkar & Orzack, 1980; Charlesworth, 1994; Wilbur & Rudolf, 2006; Morris et al., 2008; Cotto & Ronce, 2014; Ratikainen & Kokko, 2019). Using experimental evolution in the nematode *Caenorhabditis remanei*, we show that it is the temporal structure of environmental change, and not mean temperature alone, that shapes the evolution of lifespan and mortality rate. Populations in which temperature cycled every second generation (*Fast temperature cycles*) evolved a shorter lifespan and a faster increase in late-life mortality than populations held at a constant, stressful 25°C (*Warm 25°C*), which instead evolved the longest lifespan of any regime.

The traditional view has long been that variable environments select for increased somatic maintenance, so that individuals can survive unfavourable periods and reproduce when conditions improve (Murphy, 1968; Tuljapurkar & Orzack, 1980; Charlesworth, 1994; Wilbur & Rudolf, 2006; Morris *et al*., 2008). The effect is, however, expected to be small (Benton & Grant, 1999) and the theory has limited empirical support (reviewed in Roff, 2002). Because adaptation lags the environmental change (Lande & Shannon, 1996; Kopp & Matuszewski, 2014) and the strength of selection declines with age (Medawar, 1952; Hamilton, 1966; Caswell & Shyu, 2017), older individuals in an age-structured population sit furthest from the moving fitness optimum and alleles with late-acting deleterious effects therefore escape selection and ageing accelerates, following the logic of mutation accumulation theory (Medawar, 1952; Charlesworth, 1994; Cotto & Ronce, 2014). This mismatch is greatest when the environment turns over on a timescale just longer than a generation, so that individuals routinely age into environmental conditions they did not develop in, precisely the regime in which we observed the evolution of short life. Alternatively, if reversible plasticity can instead track the environment, long lifespan is expected to evolve in variable environments (Ratikainen & Kokko, 2019).

We found evidence that the populations in *Fast temperature cycles* had evolved a shorter lifespan than the *Warm 25°C* populations in both testing temperatures. In addition, they also had a higher increase in age-specific mortality rate, demonstrating that the reduced lifespan was because of faster ageing. Several lines of evidence indicate that the short lifespan of the *Fast temperature* cycles regime is adaptive. These populations are the most fit when parent and offspring temperatures are uncorrelated, as they were during evolution and in our 25°C assays, and they have lost the anticipatory maternal effect retained by the other regimes, against which it was selected because parent and offspring environments were on average uncorrelated (Lind *et al*., 2020). Critically, the absence of a selection regime × temperature interaction argues against maternal effects driving the lifespan differences between regimes, and the *Fast temperature* cycles regime was the shortest-lived in both assay temperatures. Together with the previously published fitness (Lind et al., 2020) and life-history (Sekajova et al., 2025) data, this points to a trade-off between early-life fitness under fluctuating conditions and lifespan. Also, all regimes that experienced 25°C during evolution pay a fitness cost in 20°C relative to the *Control* 20°C regime (Lind et al., 2020).

Theory have identified that the evolution of reversible plasticity in variable environments can result in the evolution of long lifespan, while evolution of irreversible developmental plasticity (or lack of plasticity) will result in phenotypic mismatch once the environment changes, and the evolution of short lifespan (Ratikainen & Kokko, 2019). Our results support these predictions. We have recently shown that the fast cycle lines have evolved increased developmental plasticity in body size as a response to environmental variation (Sekajova *et al*., 2025) which together with the short lifespan (this study) and the lack of maternal effects (Lind *et al*., 2020) fits a model of adaptation by avoiding phenotypic mismatch caused by environmental change. Irreversible plasticity is common in nematodes (Viney & Diaz, 2012) and body size is a classic example of such plasticity. Since cell division of somatic cells stops at sexual maturity in *Caenorhabditis* nematodes, there are limited possibilities for major morphological changes after maturation, except for growth of cell size.

Our second key result is that constant, stressful warmth selected for the longest lifespan of any regime, even though exposure to 25°C is itself plastically life-shortening and increases mortality rate. This appears to run counter to the prediction that maladaptation to a novel but stable environment should shorten life (Cotto & Ronce, 2014), but the two are reconciled once the selective properties of the environment itself are considered, which that general model does not capture. Warm temperature is stressful, and heat-shock resistance is repeatedly associated with long life, both within nematodes (Amrit et al., 2010; Chen & Maklakov, 2012; Lind et al., 2017) and across taxa (Rose et al., 1992; Sørensen et al., 2003; Holzenberger et al., 2003). In addition, 25°C suppresses total reproduction in *C. remanei* (Sekajova et al., 2025), and a recent model suggests that reduced reproduction alone is sufficient to result in the evolution of longer lifespan (Irish et al., 2025). Mean temperature and thermal variability thus act on the evolution of lifespan in opposite directions, where stable warmth lengthens it, while rapid fluctuation shortens it.

In conclusion, the temporal structure of environmental change, relative to generation time, is a key driver of the evolution of lifespan and mortality rate. Rapid fluctuations that leave organisms likely to age into a different environment select for short lifespan, whereas a stable but stressful warm environment selects for long lifespan. Because ongoing climate change is increasing both mean temperatures and their variability across the planet (Vasseur *et al*., 2014; Vázquez *et al*., 2017), anticipating its life-history consequences will require attention not only to how much environments change, but to how fast they change relative to the generation time of the organism.

## Acknowledgement

This work was funded by the Swedish Research Council [grant number 2016-05195 and 2020-04388 to MIL, grant number 621-2013-4828 to AAM], the European Research Council [grants AGINGSEXDIFF and GERMLINEAGEINGSOMA to AAM] and IIR was supported by the Norwegian Research Council [grant number 240008 and 223257]. We thank Frank Johansson, Anssi Laurila, and Claus Rüffler for discussions.

## Author contributions

MIL and AAM designed the experiment with aid from IIR, MIL, JA, HC, TK and TL performed experimental evolution, MIL and JA performed phenotypic assays, MIL analysed the data, MIL drafted the manuscript together with AAM and IIR. All authors contributed to the revision of the manuscript.

## References

Benton, T.G. & Grant, A. (1999) Optimal reproductive effort in stochastic, density-dependent environments. Evolution, 53, 677–688.

Botero, C.A., Weissing, F.J., Wright, J. & Rubenstein, D.R. (2015) Evolutionary tipping points in the capacity to adapt to environmental change. Proceedings of the National Academy of Sciences, 112, 184–189.

Bubliy, O.A. & Loeschcke, V. (2005) Correlated responses to selection for stress resistance and longevity in a laboratory population of *Drosophila melanogaster*. Journal of Evolutionary Biology, 18, 789–803.

Caswell, H. & Shyu, E. (2017) Senescence, selection gradients and mortality. In The evolution of senescence in the tree of life (ed. by Shefferson, R.P., Jones, O.R. & Salguero-Gómez, R.). Cambridge University Press, Cambridge, UK, pp. 56–82.

Charlesworth, B. (1994) Evolution in age-structured populations. 2nd edn. Cambridge University Press, New York, NY, USA.

Chen, H. & Maklakov, A.A. (2012) Longer life span evolves under high rates of condition-dependent mortality. Current Biology, 22, 2140–2143.

Chen, H.-y., Spagopoulou, F. & Maklakov, A.A. (2016) Evolution of male age-specific reproduction under differential risks and causes of death: males pay the cost of high female fitness. Journal of Evolutionary Biology, 29, 848–856.

Colchero, F., Jones, O.R. & Rebke, M. (2012) BaSTA: an R package for Bayesian estimation of age-specific survival from incomplete mark–recapture/recovery data with covariates. Methods in Ecology and Evolution, 3, 466–470.

Cotto, O. & Ronce, O. (2014) Maladaptation as a source of senescence in habitats variable in space and time. Evolution, 68, 2481–2493.

Hamilton, W.D. (1966) The moulding of senescence by natural selection. Journal of Theoretical Biology, 12, 12–45.

Irish, S.D., Kimberley, A., Immler, S., Moorad, J. & Maklakov, A.A. (2025) Mutation accumulation underpins evolution of lifespan extension by dietary restriction.

Karabatsos, G. (2006) Bayesian nonparametric model selection and model testing. Journal of Mathematical Psychology, 50, 123–148.

Kopp, M. & Matuszewski, S. (2014) Rapid evolution of quantitative traits: theoretical perspectives. Evolutionary Applications, 7, 169–191.

Kullback, S. & Leibler, R.A. (1951) On information and sufficiency. Annals of Mathematical Statistics, 22, 79–86.

Lande, R. & Shannon, S. (1996) The role of genetic variation in adaptation and population persistence in a changing environment. Evolution, 50, 434–437.

Lind, M.I., Chen, H., Meurling, S., Guevara Gil, A.C., Carlsson, H., Zwoinska, M.K., et al. (2017) Slow development as an evolutionary cost of long life. Functional Ecology, 31, 1252–1261.

Lind, M.I., Zwoinska, M.K., Andersson, J., Carlsson, H., Krieg, T., Larva, T., et al. (2020) Environmental variation mediates the evolution of anticipatory parental effects. Evolution Letters, 4, 371–381.

Lind, M.I., Zwoinska, M.K., Meurling, S., Carlsson, H. & Maklakov, A.A. (2016) Sex-specific trade-offs with growth and fitness following lifespan extension by rapamycin in an outcrossing nematode, Caenorhabditis remanei. Journals of Gerontology Series A: Biological Sciences and Medical Sciences, 71, 882–890.

Lionaki, E. & Tavernarakis, N. (2013) Assessing aging and senescent decline in Caenorhabditis elegans: cohort survival analysis. In Cell Senescence, Methods in Molecular Biology (ed. by Galluzzi, L., Vitale, I., Kepp, O. & Kroemer, G.). Humana Press, New York, pp. 473–484.

Medawar, P.B. (1952) An unresolved problem in biology. Lewis, London, UK.

Morris, W.F., Pfister, C.A., Tuljapurkar, S., Haridas, C.V., Boggs, C.L., Boyce, M.S., et al. (2008) Longevity can buffer plant and animal populations against changing climatic variability. Ecology, 89, 19–25.

Murphy, G.I. (1968) Pattern in life history and the environment. The American Naturalist, 102, 391–403.

Ratikainen, I.I. & Kokko, H. (2019) The coevolution of lifespan and reversible plasticity. Nature Communications, 10, 538.

Reznick, D.N., Bryant, M.J., Roff, D., Ghalambor, C.K. & Ghalambor, D.E. (2004) Effect of extrinsic mortality on the evolution of senescence in guppies. Nature, 431, 1095–1099.

Rodríguez-Muñoz, R., Boonekamp, J.J., Fisher, D., Hopwood, P. & Tregenza, T. (2019) Slower senescence in a wild insect population in years with a more female-biased sex ratio. Proceedings of the Royal Society B: Biological Sciences, 286, 20190286.

Roff, D.A. (2002) Life history evolution. Sinauer Associates Inc, Sunderland, USA.

Sekajova, Z., Fossen, E.I.F., Rosa, E., Ratikainen, I.I., Tourniaire-Blum, M., Bolund, E., et al. (2025) Evolution of phenotypic plasticity during environmental fluctuations. Journal of Evolutionary Biology, voaf078.

Sørensen, J.G., Kristensen, T.N. & Loeschcke, V. (2003) The evolutionary and ecological role of heat shock proteins. Ecology Letters, 6, 1025–1037.

Spagopoulou, F., Teplitsky, C., Chantepie, S., Lind, M.I., Gustafsson, L. & Maklakov, A.A. (2020) Silver-spoon upbringing improves early-life fitness but promotes reproductive ageing in a wild bird. Ecology Letters, 23, 994–1002.

Stearns, S.C. (1992) The evolution of life histories. Oxford University Press, New York, NY, USA.

Stearns, S.C., Ackermann, M., Doebeli, M. & Kaiser, M. (2000) Experimental evolution of aging, growth, and reproduction in fruitflies. Proceedings of the National Academy of Sciences, 97, 3309–3313.

Stiernagle, T. (2006) Maintenance of C. elegans. WormBook: the online review of C. elegans biology.

Tufto, J. (2015) Genetic evolution, plasticity, and bet-hedging as adaptive responses to temporally autocorrelated fluctuating selection: A quantitative genetic model. Evolution, 69, 2034–2049.

Tuljapurkar, S.D. & Orzack, S.H. (1980) Population dynamics in variable environments I. Long-run growth rates and extinction. Theoretical Population Biology, 18, 314–342.

Vasseur, D.A., DeLong, J.P., Gilbert, B., Greig, H.S., Harley, C.D.G., McCann, K.S., et al. (2014) Increased temperature variation poses a greater risk to species than climate warming. Proceedings of the Royal Society B: Biological Sciences, 281, 20132612.

Vázquez, D.P., Gianoli, E., Morris, W.F. & Bozinovic, F. (2017) Ecological and evolutionary impacts of changing climatic variability. Biological Reviews, 92, 22–42.

Viney, M. & Diaz, A. (2012) Phenotypic plasticity in nematodes: Evolutionary and ecological significance. Worm, 1, 98–106.

Wilbur, H.M. & Rudolf, V.H.W. (2006) Life-history evolution in uncertain environments: bet hedging in time. The American Naturalist, 168, 398–411.

Williams, P.D., Day, T., Fletcher, Q. & Rowe, L. (2006) The shaping of senescence in the wild. Trends in Ecology & Evolution, 21, 458–463.

