## Supplementary table 1,2 and 3, supplementary figure 1 for "Rapid environmental change favours the evolution of shorter lifespan"

### Variable environments select for short lifespan

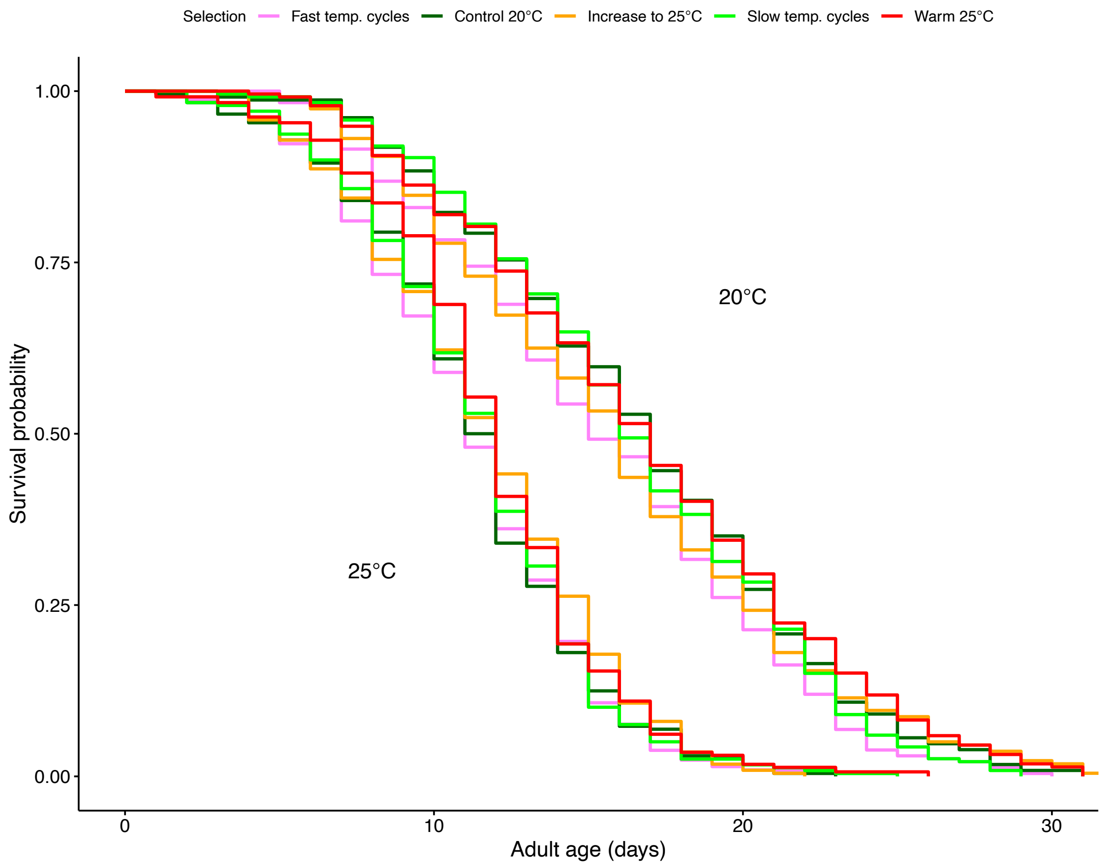


**Supplementary figure 1.** Survival curves in 20°C and 25°C. Line colour represent experimental evolution regime.

**Supplementary table 1.** Pre-planned post-hoc contrasts of survival between experimental evolution regimes, averaged across temperatures.

| Contrast | Estimate | SE | Z-ratio | p-value |
|---|---|---|---|---|
| Control 20°C – Fast temp. cycles | -0.125 | 0.0714 | -1.752 | 0.241 |
| Increased warming - Fast temp. cycles | -0.159 | 0.0719 | -2.219 | 0.090 |
| Slow temp. cycles - Fast temp. cycles | -0.111 | 0.0711 | -1.565 | 0.334 |
| Warm 25°C - Fast temp. cycles | -0.219 | 0.0720 | -3.043 | 0.009 |

**Supplementary table 2.** All possible post-hoc contrasts of survival between experimental evolution regimes, averaged across temperatures.

| Contrast | Estimate | SE | Z-ratio | p-value |
|---|---|---|---|---|
| Fast temp. cycles - Control 20°C | 0.125 | 0.0714 | 1.752 | 0.402 |
| Fast temp. cycles - Increased warming | 0.159 | 0.0719 | 2.219 | 0.173 |
| Fast temp. cycles - Slow temp. cycles | 0.111 | 0.0711 | 1.565 | 0.520 |
| Fast temp. cycles – Warm 25°C | 0.219 | 0.0720 | 3.043 | 0.020 |
| Control 20°C - Increased warming | 0.034 | 0.0715 | 0.481 | 0.989 |
| Control 20°C – Slow temp. cycles | -0.014 | 0.0709 | -0.193 | 1.000 |
| Control 20°C – Warm 25°C | 0.094 | 0.0716 | 1.312 | 0.684 |
| Increased warming – Slow temp. cycles | -0.048 | 0.0714 | -0.674 | 0.962 |
| Increased warming – Warm 25°C | 0.060 | 0.0719 | 0.827 | 0.922 |
| Slow temp. cycles – Warm 25°C | 0.108 | 0.0714 | 1.506 | 0.559 |

**Supplementary table 3.** KLDC values for all mortality parameter comparisons. KLDC-values larger than 0.8 indicate significantly different distributions.

| Comparison | c | $b_0$ | $b_1$ |
|---|---|---|---|
| 25°C - 20°C | 0.961 | 0.959 | 0.992 |
| Increased warming - Fast temp. cycles | 0.671 | 0.609 | 0.796 |
| Slow temp. cycles - Fast temp. cycles | 0.628 | 0.945 | 0.884 |
| Warm 25°C - Fast temp. cycles | 0.587 | 0.557 | 0.948 |
| Slow temp. cycles – Increased warming | 0.510 | 0.980 | 0.531 |
| Warm 25°C - Increased warming | 0.534 | 0.765 | 0.638 |
| Warm 25°C - Slow temp. cycles | 0.508 | 0.872 | 0.545 |
